# Inhibitory GABAergic neuron loss due to oxidative damage during *ex vivo* acute brain slice preparation influences genesis and dynamics of epileptiform activity

**DOI:** 10.1101/2025.10.08.681234

**Authors:** Felix Chan, Anupam Hazra, Ashan Jayasekera, Katherine Huang, Shuna Whyte, Leolie Telford-Cooke, Kamilah Lakhani, Xiaomeng Li, Rebecca Shields, Angeline Kosim, Darwin Su, Carol Murray, Mark O. Cunningham

## Abstract

*Ex vivo* acute brain slice is a popular technique in neuroscience research. Since its inception five decades ago, many variations of the brain slice preparation method have emerged. While all variations are currently used by many labs throughout the world, no study has comprehensively examined the impact of these variation on the quality of the acute brain slice preparation. In this study, we comprehensively examined the effect of animal sacrifice methods (decapitation or transcardial perfusion) and cutting solution (normal or sucrose artificial cerebrospinal fluid) on brain slice preparation. Neuronal population was quantified by immunohistochemistry against various neuronal markers. Neuronal dynamics was evaluated by *in vitro* electrophysiology using two acute epilepsy models – zero-magnesium and 4-aminopyridine. To modulate the stress incurred in acute brain slice preparation, we administered antioxidants or HIF-1α inhibitor, chrysin, to the cutting solution. The method of brain slice preparation significantly affected the quality of the brain slice preparation. In general, the combination of transcardial perfusion and sucrose artificial cerebrospinal fluid produces the optimal brain slice preparation. This is evidenced by a preservation of inhibitory GAαBAergic neurons in slices prepared with this combination. We subsequently found that this loss of inhibitory GABAergic neurons significantly influenced the genesis and dynamics of induced acute epileptiform activity. The slices with preserved inhibition had less successful induction of acute epileptiform activity and a seizure-like event that is typical of those induced in brain with preserved inhibition. Finally, we found that loss of inhibitory GABAergic neurons during brain slice preparation is primarily due to oxidative damage. Limiting oxidative stress is an effective neuroprotection strategy to prevent loss of inhibition in brain slice preparation. In conclusion, consideration of brain slice preparation method is crucial in preserving inhibitory GABAergic neurons and the degree of inhibition in *ex vivo* acute brain slice preparation.

## Introduction

Since its inception in 1957 by Li and McIlwain (Li & McIlwain, 1957), *ex vivo* acute brain slice electrophysiology has proven to be a very popular technique in interrogating the functional state of brain activity. Many modifications to this technique have existed since then – one of import is the use of sucrose artificial cerebrospinal fluid (sACSF) to reduce excitotoxicity (Aghajanian & Rasmussen, 1989). This study had reported no viable motoneuron in slices prepared using normal artificial cerebrospinal fluid (nACSF) and strikingly, a preservation of viable motoneuron in sACSF-prepared slices (Aghajanian & Rasmussen, 1989). Since then, several other studies have reported differences in the functional state of brain slice prepared in different solution, including significant change in long-term potentiation (LTP) induction and gamma-aminobutyric acid(GABA)-ergic neurotransmission (Avegno et al., 2019; Kuenzi et al., 2000). Collectively, it can be said that the composition of the ACSF as the cutting solution in brain slice preparation directly influence neuronal survival and function; and ultimately the quality of the brain slice preparation (Richerson & Messer, 1995).

Another major adaptation in the slice preparation technique is the method of animal sacrifice and brain extraction. Classically, animals such as rodent were simply decapitated with or without anesthetic agents and brain was obtained speedily from this preparation. Indeed, this method is still used presently and can still be seen in published standard operating protocol for brain slice preparation (Papouin & Haydon, 2018). Another popular method of euthanasia is transcardial perfusion of rodents. Typically used to perfuse fixative to preserve brain tissue for histology (Wu et al., 2021), transcardial perfusion of ACSF, either nACSF or a modified ACSF such as sACSF and others, has been used to prepare acute brain slice for electrophysiology purposes (Avegno et al., 2019; Chan et al., 2019). Many other variations to this technique exist, such as anaesthetic agent used (Schurr et al., 1995), temperature of slicing, and slicing techniques; and while attempt has been made at creating a consensus and standard of practice for *ex vivo* brain slice preparation (Raimondo et al., 2017), individual research groups still maintain their own standard operating procedure for brain slice electrophysiology.

Epilepsy is a major area of neuroscience research where *ex vivo* acute brain slice electrophysiology is commonly used. A recent report from the International League Against Epilepsy Task Force clearly recognized the importance of using acute brain slice preparation as a model for epilepsy study, particularly for high throughput screening and therapeutic development (Morris et al., 2023). However, as mentioned previously, the use of acute brain slice model of epilepsy varies by method of induction, variations in the protocol, and experimentation in specific research groups (Raimondo et al., 2017). The different acute model of epilepsy targets different method of induction; such as increasing N-methyl-D-aspartate receptor (NMDA-R) mediated excitation using the zero magnesium (0 Mg^2+^) model (Mody et al., 1987), increasing potassium-channel activity using 4-aminopyridine (Brückner & Heinemann, 2000), or reducing GABAergic inhibition using GABA_A_ receptor antagonists (for e.g., picrotoxin, pentylenetetrazole or bicuculline (Hashimoto et al., 2017; Müller et al., 2018; Samoilova et al., 2003). Despite the well-established model of epilepsy in acute brain slice preparation, no study has directly evaluated the influence of different method of acute brain slice preparation on the induction and dynamics of epileptiform activity. This is particularly surprising given that as mentioned in aforementioned studies, change in method of brain slice preparation can have dramatic effect on synaptic properties, inhibition, and neuronal survival (Aghajanian & Rasmussen, 1989; Avegno et al., 2019; Kuenzi et al., 2000; Richerson & Messer, 1995)all of which can affect the epileptiform activities induced.

This study fills in this important gap in our knowledge by comprehensively examining the impact of different brain slice preparation technique on the dynamic and induction of epileptiform activity. We will focus on two aspects of brain slice preparation technique which we hypothesize pose the most significant determinant factor: (1) the composition of the cutting solution; either nACSF or sACSF and (2) transcardial perfusion vs decapitation. Of interest to us is specifically the effect of this variation in brain slice preparation technique on the inhibitory interneuron population in the brain. Inhibitory interneurons are crucial in shaping the epileptic network dynamics (Muldoon et al., 2015). Additionally, due to its high firing rate, they often have higher metabolic demand and are thus, more vulnerable to oxidative damage (Kageyama & Wong-Riley, 1982; Wang & Michaelis, 2010). Thus, we hypothesize that these interneuron population may be the most affected by the difference in slice preparation technique and ultimately, influence the outcome of the epileptiform activity induction.

## Methods

### Ethical approval and use of animals

Adult (10-12 weeks old) male Wistar rats were used in this study. All animal handling and experimentation were done in accordance with the requirements of the United Kingdom Animals (Scientific Procedures) Act 1986. Animals were housed on a 12 h light-dark cycle under controlled conditions (temperature: 20°C–25°C; humidity: 40%–60%). Food and water were available ad libitum.

### Brain slice preparation

Brain slices used for the *in vitro* studies were prepared using either one of the following protocols. In the first paradigm; which we termed ‘decapitated’, the animals were exposed to inhaled isoflurane (IsoFlo®, Abbott Laboratories Ltd) and after deep anaesthesia was achieved, cervical dislocation was performed. The rats were then decapitated and brains were rapidly removed in cutting solution. In the second paradigm, which we termed ‘perfused’, the animals were again exposed to inhaled isoflurane and terminal anaesthesia was given by injection of 30mg of ketamine (Narketan®, Vetoquinol) and 6mg of xylazine (Xylacare®, Animalcare). After all reflexes were lost, we performed transcardial perfusion of the cutting solution. The brain was then rapidly removed, again, in the cutting solution.

Two cutting solutions were tested in this study, namely sACSF; which contains 252mM sucrose, 24mM NaHCO_3_, 2mM MgSO_4_, 2mM CaCl_2_, 3.5mM KCl, 1.25mM NaH_2_PO_4_, and 10mM glucose; and nACSF; which contains 126mM NaCl, 24 mM NaHCO_3_, 1.2mM MgSO_4_, 1.2mM CaCl_2_, 3mM KCl, 1.25mM NaH_2_PO_4_, and 10mM glucose. All the cutting solutions were prepared ice-cold and oxygenated (95% O_2_/5% CO_2_).

By combining two slice preparation methods with two different cutting solutions, we systematically tested four variations of brain slice preparation techniques, namely (1) decapitated with sACSF (DS), (2) decapitated with nACSF (DN), (3) perfused with sACSF (PS), and (4) perfused with nACSF (PN). The remaining slice preparation protocol is the same for any of the four paradigms. The brains were sliced using a vibratome (5100mz, Camden Instruments) in the transverse orientation (450µm thickness). Slices containing the entorhinal cortex were collected and placed in an interface holding chamber in oxygenated nACSF at room temperature for one hour before any experimentation.

For the rescue studies, two additional cutting solutions were utilised: (1) a nACSF solution containing a cocktail of antioxidants (10mM ascorbic acid, 100µM alpha-tocopherol, and 2mM N-acetylcysteine) (resulting in a paradigm called DA – when combined with decapitation) and (2) a nACSF solution containing chrysin (0.05% v/v) – called chrysin cutting solution (resulting in a paradigm called DC – when combined with decapitation).

### Electrophysiology

For local field potential recordings, brain slices were transferred to interface recording chamber maintained at 30-33^0^C which is continuously perfused with oxygenated nACSF solution. Slices were left to incubate in the recording chamber for 30 minutes. To evoke epileptiform activity, we used two commonly studied *in vitro* model of seizures: (1) zero Mg^2+^ model (where a Mg^2+^-free aCSF formulation is used) and 4-aminopyridine (4-AP; 200µM) model. Local field potentials were recorded with aCSF filled microelectrodes positioned in the layer 2-3 of the medial entorhinal cortex (mEC). The signal was amplified using the EXT10-2F differential amplifier (NPI Electronic), filtered online (0.1-500Hz), passed through a 50Hz noise eliminator (Hum-Bug, Quest Scientific), digitized by ITC-18 acquisition board (Instrutech), sampled at 5kHz, and acquired using Axograph X (Axograph Scientific) software. All electrophysiological recordings were analysed using AxoGraph X software by an experimenter blinded to the experimental conditions.

### Immunohistochemistry

For immunohistochemistry, brain slices were prepared according to the different slice preparation paradigm as described above and incubated in the holding chamber for 3 hours. Following this, the slices were fixed in 4% paraformaldehyde in 0.1M phosphate-buffered saline and stored at 4^0^C until processing. Brain slice processing and immunohistochemistry was conducted using previously published protocol (Chan et al.,2019). The primary and secondary antibodies used in this study were listed in Supplementary Table 1. Sections were visualized using a stereology light microscope (Olympus, BX51) and the software StereoInvestigator (MBF Bioscience) was used to perform the cell count. A region was drawn on the medial entorhinal cortex layers I, II, and III of each brain sections and counts were normalized by the area of the region to give cell density (cells/mm^2^) values.

### Biochemical assay

For the biochemical assay, brain slices were prepared according to the different slice preparation paradigm as described above and incubated in the holding chamber for 3 hours. Four brain slices were homogenized into one pooled sample using the T-18 digital Ultra-Turrax blender (IKA Labortechnik) through a 3x15 seconds blend cycle, with 30 seconds break in-between, in 1ml of ice-cold MES buffer (50mM MES, 1mM EDTA, 1% Tween-20, 1mM PMSF, and 1x protease inhibitor cocktail). The resulting homogenate was then centrifuged at 10,000xg for 10 minutes at 4^0^C to separate the insoluble fraction. The supernatant was extracted and snap-frozen at -80^0^C until processed. Carbonyl detection was performed as per established method on derivatization with 2,4-dinitrophenylhydrazine (DNPH) and detected through colorimetric assay (Levine et al., 1990). The absorbance was read at 370nm using the SpectraMax M3 plate reader (Molecular Devices) and was corrected by subtracting the control absorbance from the sample absorbance for each sample. The corrected protein carbonyl value was then normalized against the protein concentration as measured by a standard Bradford assay.

### Statistical analysis

Data are displayed as mean ± SEM. Normality of the dataset was tested using the Shapiro-Wilk test. Statistical significance testing for normally distributed datasets was conducted using either a t-test or ANOVA (one-factor or two-factor) with post-hoc t-test and correcting for multiple comparisons using Prism (GraphPad). In cases of datasets that were not normally distributed, equivalent non-parametric tests were utilized. Statistical significance was accepted at *P*<0.05.

## Results

### Neuronal viability varies with brain slice preparation method

We comprehensively assessed the viability of neurons in different methods of brain slice preparations (see Figure 1) firstly by detecting the expression of constitutively expressed nuclear protein, NeuN. Brain slices prepared using perfusion with sucrose aCSF (PS) (1,115.0 ± 46.5 cells/mm^2^) had the largest number of NeuN-positive cells and this was significantly greater (p<0.05) than brain slices prepared using decapitation with normal ACSF (DN) (738.5 ± 58.1 cells/mm^2^) or decapitation with sucrose ACSF (DS) (850.1 ±99.9 cells/mm^2^). This significant reduction in NeuN expression indicated that there was a significant loss of neurons in the brain slices prepared using the decapitation method. As NeuN is expressed in both glutamatergic and GABAergic neurons, we next examined,using antibodies to Ca ^2+^/calmodulin-dependent protein kinase II (CaMKII) and GABA, if the observed reduction in neuronal population was specific to excitatory or inhibitory neurones respectively. Brain slices prepared using the PS method (423.5 ± 48.4 cells/mm^2^) exhibited the largest immunoreactivity for GABA and this was significantly greater than any of the other methods (DN; 152.3 ± 23.7 cells/mm^2^, DS; 155.8 ± 18.84 cells/mm^2^, PN; 236.2 ± 36.8 cells/mm^2^). Similarly, the largest immunoreactivity for CaMKII was observed in brain slices prepared using the PS method (285.6 ± 42.7 cells/mm^2^) and this was significantly greater than slices prepared using DN (137.1 ± 20.9 cells/mm^2^) or DS (137.1 ± 18.7 cells/mm^2^) method.

**Figure 1:**
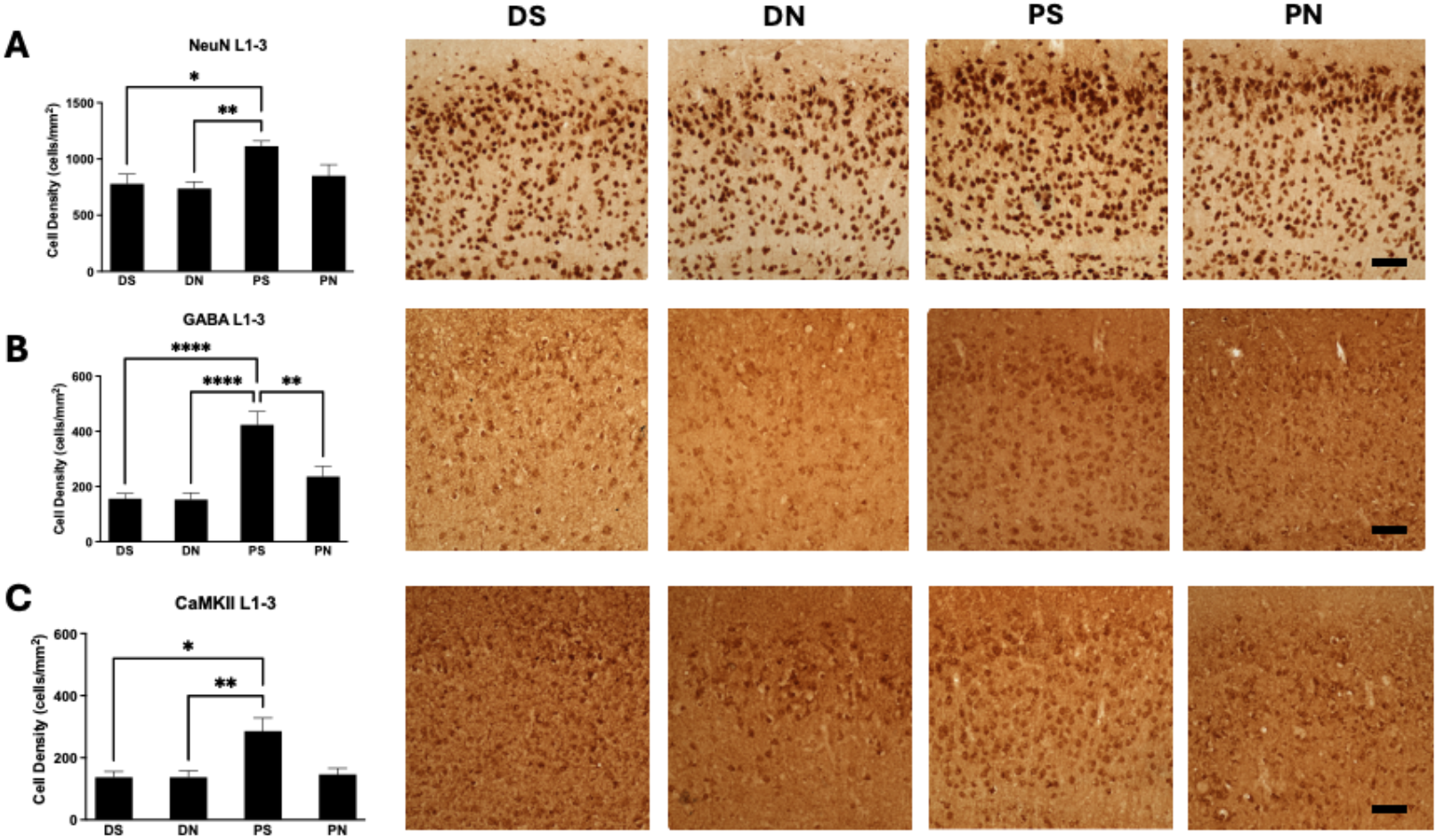
Differing slice preparation methods alter neuronal population *in vitro*. In each panel is shown representative photomicrographs of the staining profile for NeuN (A), GABA (B), and CaMKII (C) respectively. Alongside the photomicrographs, a bar chart quantifying the cell density of each slice preparation groups is shown with statistics. DS – decapitated with sucrose aCSF, DN – decapitated with normal aCSF, PS – perfused with sucrose aCSF, PN – perfused with normal aCSF. * p<0.05, ** p<0.01, *** p<0.001, **** p<0.0001. n for DS – 6, DN – 8, PS – 7, PN – 8. Scale bar – 10µm.

**Figure 2:**
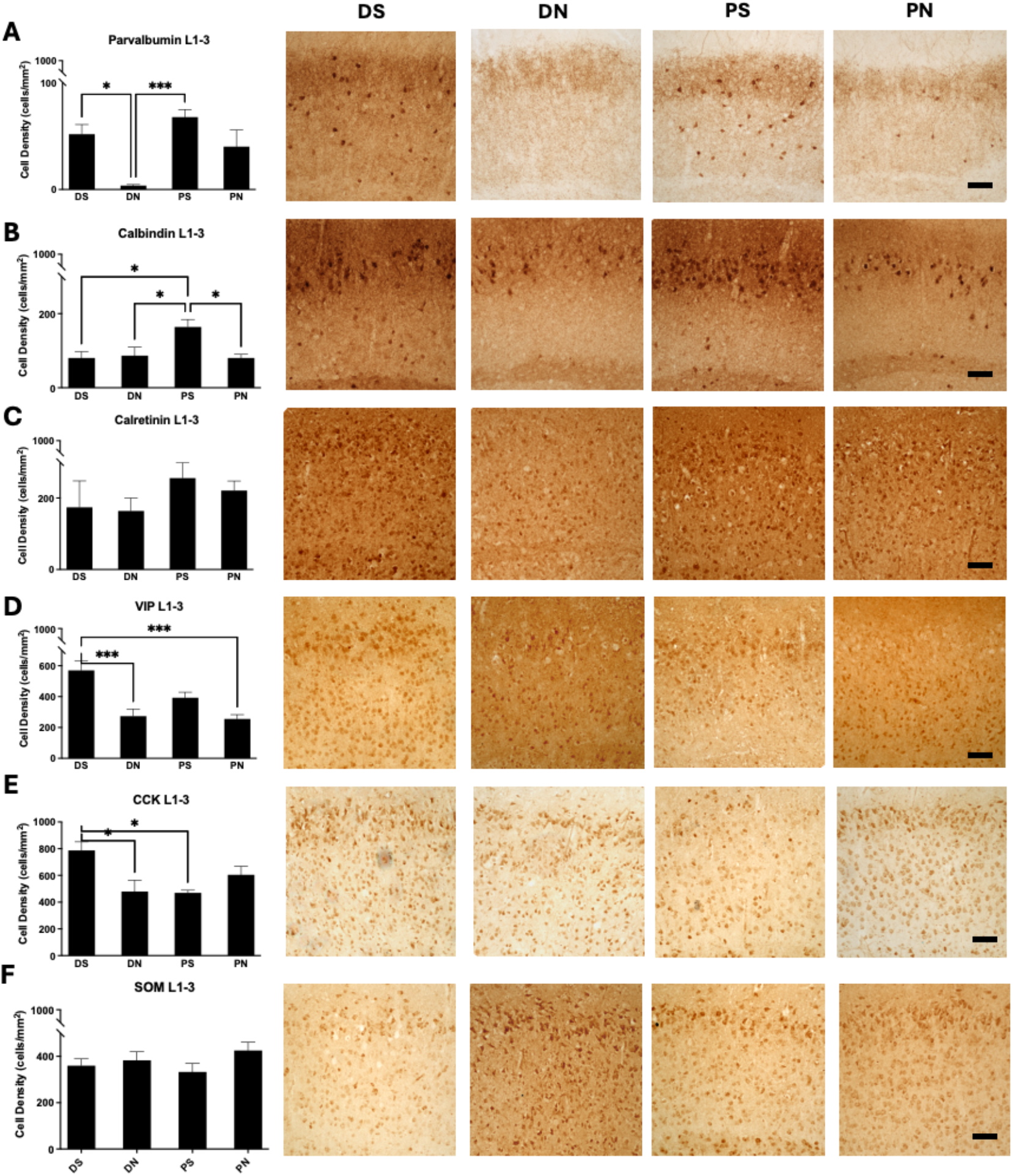
Inhibitory interneuron subtypes are differentially affected by variations in slice preparation methods. In each panel is shown representative photomicrographs of the staining profile for parvalbumin (A), calbindin (B), calretinin (C), VIP (D), CCK (E), and somatostatin (F) respectively. Alongside the photomicrographs, a bar chart quantifying the cell density of each slice preparation groups is shown with statistics. DS – decapitated with sucrose aCSF, DN – decapitated with normal aCSF, PS – perfused with sucrose aCSF, PN – perfused with normal aCSF. * p<0.05, ** p<0.01, *** p<0.001. n for DS – 8, DN – 7, PS – 12, PN – 6. Scale bar – 10µm.

### Specific inhibitory interneuron subtypes are selectively lost across different brain slice preparation methods

There is a high degree of heterogeneity in terms of the physiology, anatomy, and molecular phenotype of cortical interneurons (Lim et al., 2018). Results in the previous section demonstrated a profound difference in the number of inhibitory GABAergic interneurons in acute brain slices depending on the slice preparation methods. We next aimed to examine if a specific subtype of interneuron is preferentially affected by the difference in the method of brain slice preparation. This is done by examining the immunoreactivity for various calcium binding proteins that serve as biomarker for discrete populations of inhibitory interneurons.

Interestingly, we found that parvalbumin-expressing (PV^+^) interneurons are significantly affected by slice preparation methods with brain slices prepared using the DN method demonstrating the lowest density of PV^+^ interneurons (DN; 3.7 ± 1.0 cells/mm^2^) which was significantly lower than slices prepared using the DS or PS method (DS; 51.6 ± 9.0 cells/mm^2^, PS; 67.6 ± 7.0 cells/mm^2^). Slices prepared using PS method showed the highest density of calbindin-expressing interneurons (PS; 164.0 ± 20.3 cells/mm^2^) which was significantly higher than slices prepared with any other preparation methods (DN; 86.7 ± 22.6 cells/mm^2^, DS; 80.3 ± 17.2 cells/mm^2^, PN; 80.5 ± 9.9 cells/mm^2^). Slices prepared using DS method showed enrichment of both vasoactive intestinal peptide (VIP^+^)- and cholecystokinin (CCK^+^)-expressing interneurons. Specifically, the VIP^+^ interneurons in DS-prepared slices (DS; 392.6 ± 35.8 cells/mm^2^) was significantly higher than DN- and PN-prepared slices (DN; 273.9 ± 45.4 cells/mm^2^, PN; 255.7 ± 27.8 cells/mm^2^). The CCK^+^ interneurons in DS-prepared slices (DS; 787.3 ± 63.6 cells/mm^2^) were significantly higher than DN- and PS-prepared slices (DN; 479.5 ± 84.6 cells/mm^2^, PS; 469.9 ± 19.9 cells/mm^2^). There was no significant difference in the expression of calretinin (CR^+^)-expressing or somatostatin (SOM^+^)-expressing interneurons among the different brain slice preparation methods.

### Interneuron preservation across slice preparation methods affects network excitability and function

Given the significant differences in the level of inhibitory and excitatory neuron composition following different brain slice preparation methods, we hypothesized that this would result in a difference in the network excitability and function. We tested this using an acute epileptiform induction model - the 0 Mg^2+^ model, a widely established model to study seizure initiation and dynamics *in vitro* (Anderson et al., 1986). The 0 Mg^2+^ model induces epileptiform activity by increasing excitability via excessive activation of the NMDAR following removal of the voltage-dependent block mediated by the Mg^2+^ ions (Fekete & Wang, 2025). As different brain slice preparation method influences excitatory-inhibitory balance, we speculated that the probability of seizure induction and its dynamic would be affected by the variations in this methodology.

The 0 Mg^2+^ model induces two form of activity – in the early phase, seizure like event (SLE) are recorded (Jones, 1989; Jones et al., 1992); and after some regular occurrences of these SLEs, the network transitioned towards late recurrent discharges (LRDs) (Li Zhang et al., 1995) – see Figure 3A. Interestingly, variations in the preparation methodology produced a significant difference in the proportion of slices displaying SLE with slices prepared using DN method more likely to experience SLE in the 0 Mg^2+^ model, *x*^2^ (3, N= 70)=18.2; p=0.0004. However, although the SLE induction probability in DS, PS, or PN are similar, the dynamics of the SLEs induced vary. The time to induce SLEs is significantly lower in slices prepared using DN method (DN; 9.5 ± 2.3 minutes) compared against DS or PS methods (DS; 46.8 ± 6.4 minutes, PS; 42.9 ± 8.0 minutes). The duration of SLEs is also significantly lower in slices prepared using DN method (DN; 12.2 ± 1.9 seconds) compared against DS or PS methods (DS; 56.0 ± 16.2 seconds, PS; 48.2 ±10.3 seconds). SLE dynamics did not differ significantly between the DN and PN methods, suggesting that the cutting solution composition has a greater impact on SLE induction.

**Figure 3:**
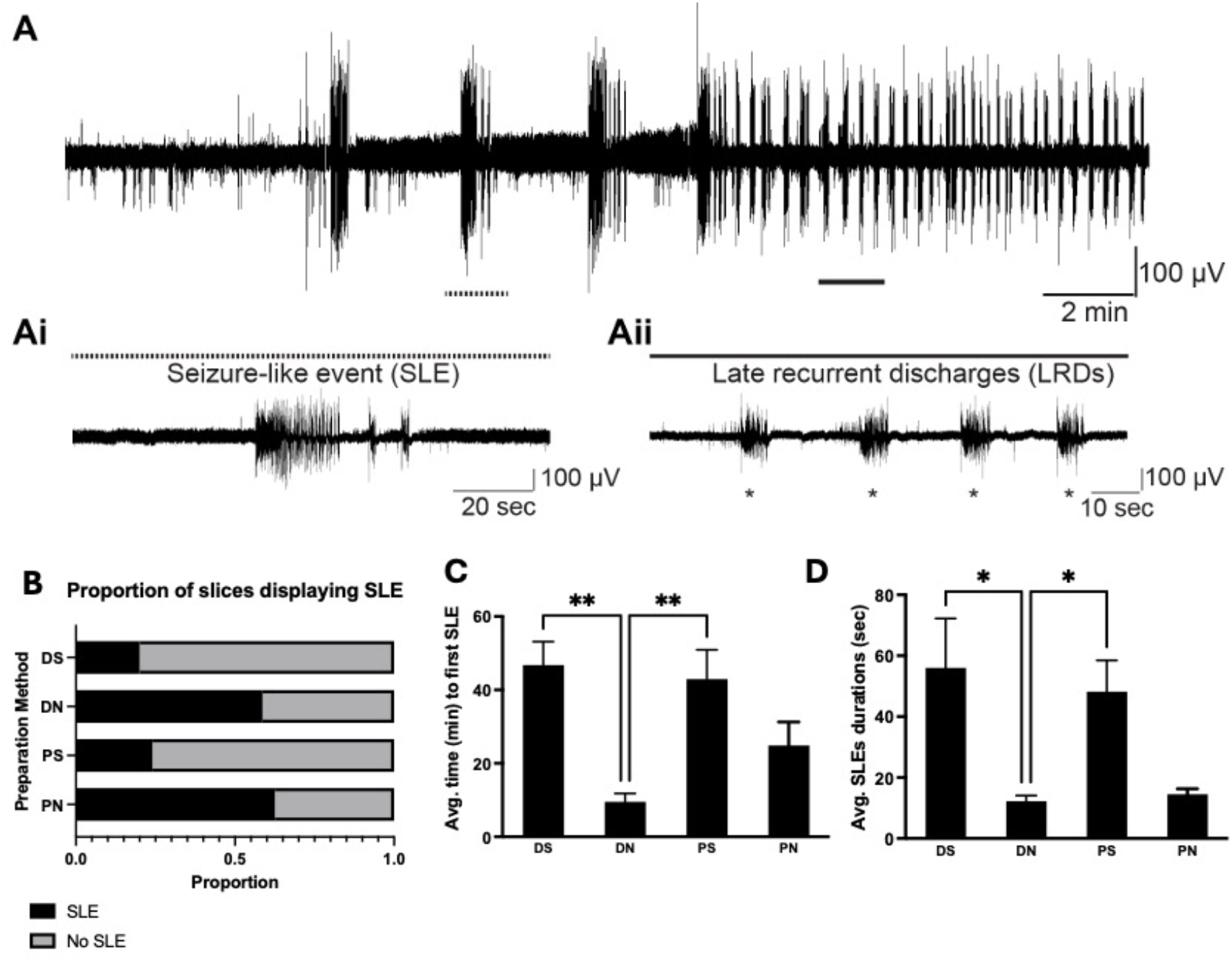
Seizure-like event (SLE) induction is significantly affected by variation in brain slice preparation method in the 0 Mg^2+^ model. In panel A is shown an example trace showing typical early induction of SLEs – zoomed in at Ai – and transitioning to late recurrent discharges (LRDs) – zoomed in at Aii. Shown in B is proportion of slices displaying SLE arranged by different brain slice preparation methods. Shown in C and D are the average time to first SLE (C) in minutes and average SLE durations (D) in seconds. DS – decapitated with sucrose aCSF, DN – decapitated with normal aCSF, PS - perfused with sucrose aCSF, PN – perfused with normal aCSF. * p<0.05, ** p<0.01, n’s for DS: 6, DN: 7, PS: 7, PN: 9.

To ensure that the observed change in network activity is not specific to the 0 Mg^2+^ model, we also tested the use of 4-AP acute epileptiform induction model in the various brain slice preparation methods. 4-AP blocks voltage-gated potassium channel, which reliably induces seizure-like events (SLEs) in acute brain slice preparation (Barbarosie & Avoli, 1997; Barbarosie et al., 2002; Heuzeroth et al., 2019). 4 -AP induces SLEs similar to those in the zero-magnesium model, but via a distinct induction mechanisms.

The 4-AP model induces two forms of activity: inter-ictal bursts and seizure-like event (SLE) similar in nature to the SLE in the 0 Mg^2+^ model – see Figure 4A. Given the similarity of the SLE in 4-AP model to the SLE in 0 Mg^2+^ model, we chose to analyze and quantify the parameters of the SLEs induced by 4-AP in the different brain slice preparation methods. Indeed, variation in brain slice preparation method similarly causes a significant difference in the proportion of slices displaying SLE with slices prepared using DN method always consistently experiencing SLE on the 4-AP model – unlike all the other slice preparation methods, *x*^2^ (3, N= 94)=34.1; p<0.0001. Interestingly, the time to induce SLEs is not significantly different between the different brain slice preparation methods, although slices prepared using DN methods showed a trend towards an earlier SLE induction time, as in the 0 Mg^2+^ model. The duration of SLEs is significantly affected with slices prepared using DN method (DN; 29.1 ± 4.9 seconds) and PN method (PN; 14.5 ±1.8 seconds) exhibiting shorter duration of SLEs compared against their sACSF prepared counterparts (DS; 73.7 ± 8.7 seconds, PS; 49.9 ± 10.4 seconds). Regardless of the epilepsy induction model, the method of brain slice preparation significantly influenced both the likelihood of successful SLE induction and the dynamics of the resulting events.

**Figure 4:**
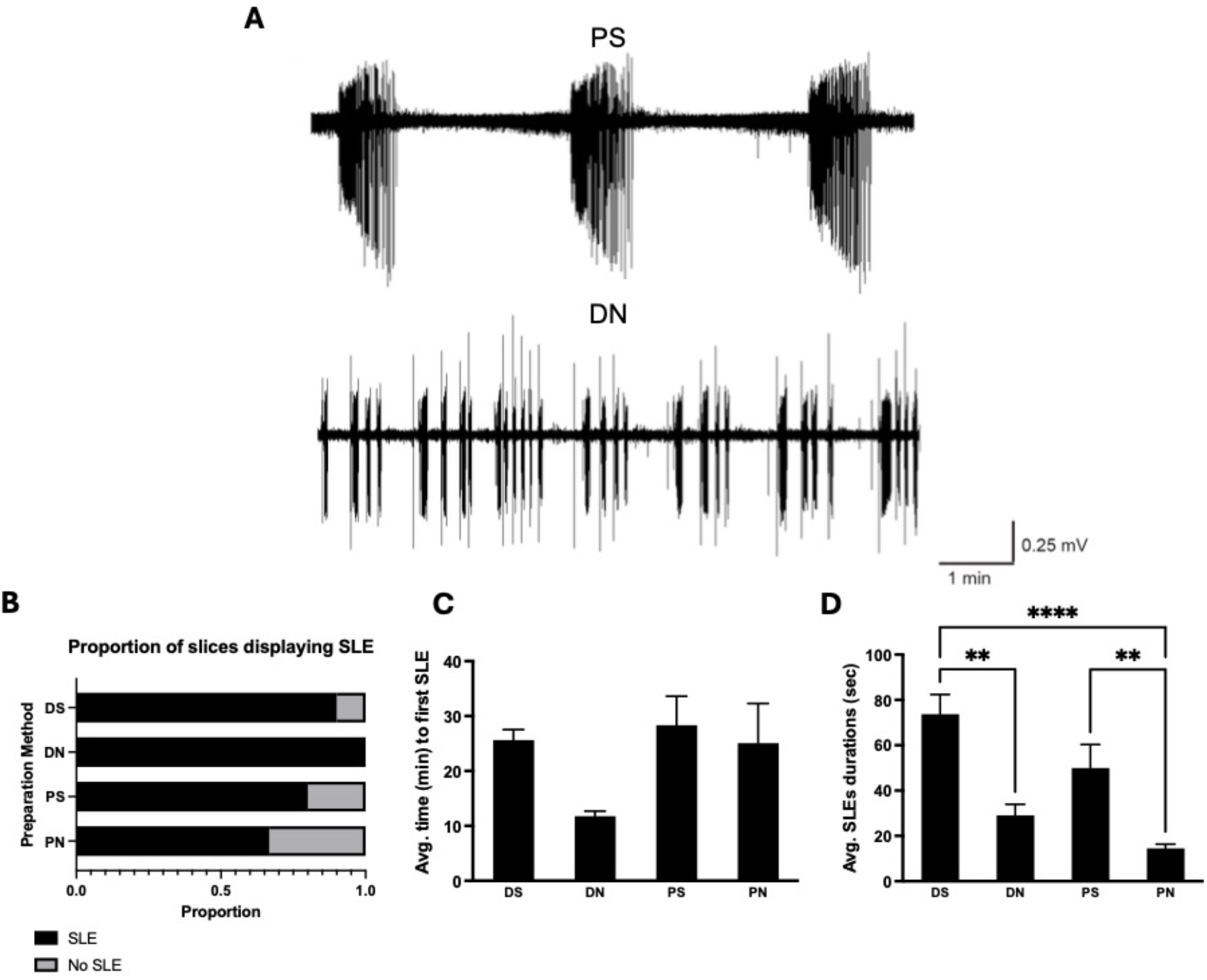
Seizure-like event (SLE) induction is significantly affected by variation in brain slice preparation method in the 4-AP model. In panel A is shown an example trace showing typical traces from 4-AP induction with differences in epileptiform activity shown in PS and DN. Shown in B is proportion of slices displaying SLE arranged by different brain slice preparation methods. Shown in C and D are the average time to first SLE (C) in minutes and average SLE durations (D) in seconds. DS – decapitated with sucrose aCSF, DN – decapitated with normal aCSF, PS – perfused with sucrose aCSF, PN – perfused with normal aCSF. ** p<0.01, **** p<0.0001, n for DS – 7, DN – 7, PS – 8, PN – 8.

### Limiting oxidative damage during brain slice preparation can preserve inhibitory interneurons

Due to their intrinsically high firing rates (in some cases > 100 Hz), inhibitory interneurons impose significant metabolic demands on the brain. (Kageyama & Wong-Riley, 1982; Wang & Michaelis, 2010). This leads to inhibitory interneurons being particularly vulnerable to oxidative damage (Wang & Michaelis, 2010), a natural byproduct of high levels of oxidative metabolism. To rescue the loss of inhibitory interneurons, we supplemented the DN cutting solution with potent antioxidant combinations, namely ascorbic acid (Gęgotek & Skrzydlewska, 2022), α-tocopherol (Robeson & Baxter, 1943), and N-acetylcysteine (Ezeriņa et al., 2018). These antioxidant combinations were chosen as they have been successfully incorporated into cutting solution compositions that were formulated specifically for preserving *ex vivo* human brain slices (Jones et al., 2016; Ting et al., 2018).

Addition of antioxidants in the DN cutting solution (see Figure 5) significantly affected the proportion of slices displaying SLEs, even if they are prepared through decapitation, *x*^2^ (1, N= 21)=7.8; p=0.005. Specifically, the proportion of slices displaying SLEs in decapitated mice prepared in nACSF cutting solution containing antioxidants (28.6%) were significantly lower than mice prepared in nACSF cutting solution (58.6%). Interestingly, there is no significant difference in the average time to first SLE. However, there is significant increase in the duration of SLEs from 12.2 ± 1.9 seconds to 21.8 ± 1.4 seconds, an SLE with duration more typical of the other slice preparation methods. We measured carbonyl concentrations as measure of protein peroxidation (Levine et al., 1990), a readout of oxidative stress. As predicted, carbonyl concentration in the brain slices is reduced in the antioxidant-treated brain slices, although this did not reach statistical significance (p=0.07). Finally, we examined the density of PV^+^ interneurons, which were significantly loss during the DN preparation. Interestingly, supplementation of antioxidants significantly rescued the interneuron loss, increasing its cell density (DA, 18.7 ± 4.5 cells/mm^2^; DN, 3.7 ± 1.0 cells/mm^2^).

**Figure 5:**
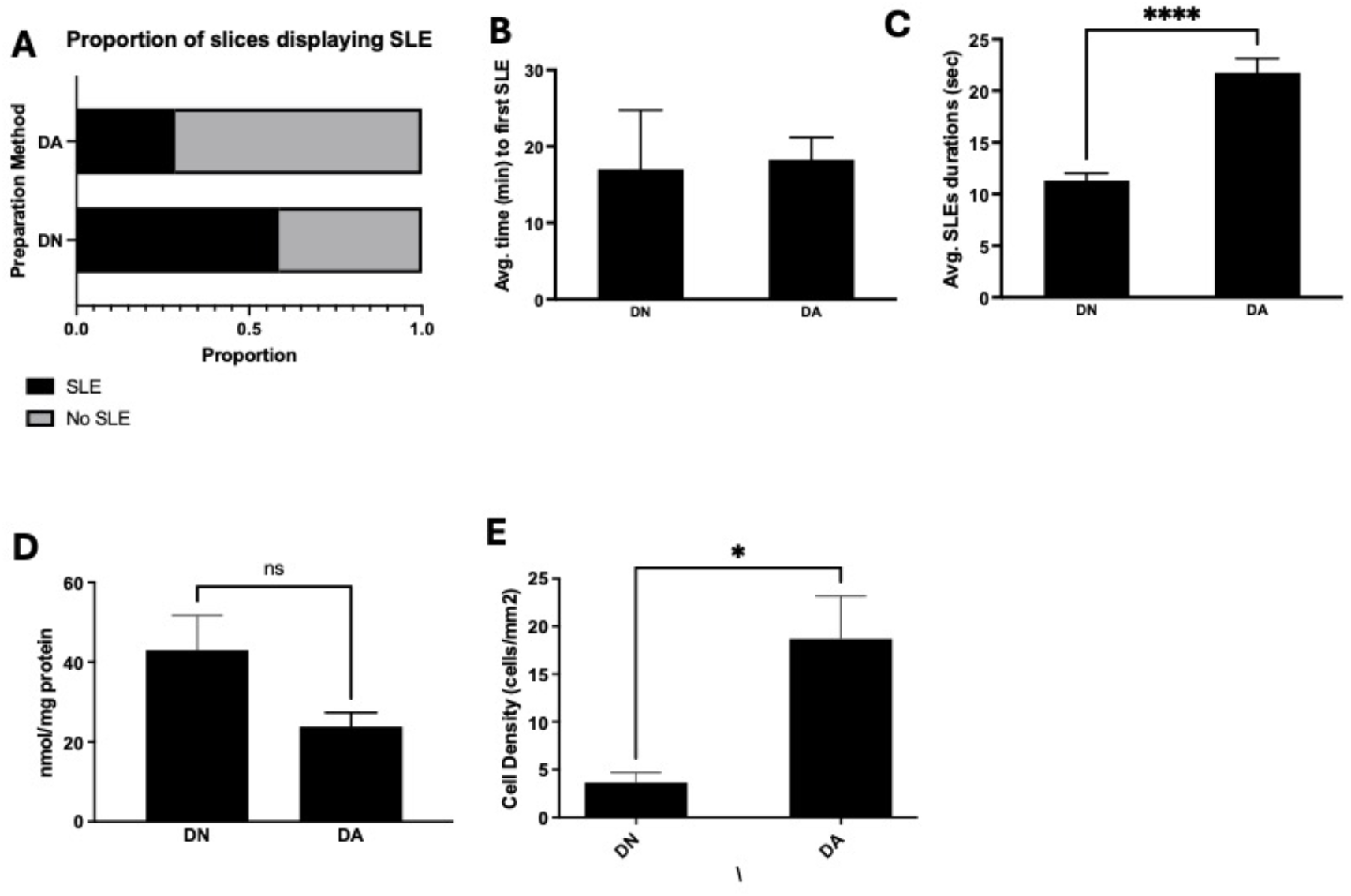
Seizure-like event (SLE) induction is rescued by addition of antioxidants in the cutting solution. In panel A is shown the proportion of slices displaying SLE in mice prepared with decapitation in nACSF cutting solution supplemented with antioxidants (DA) compared against nACSF only cutting solution. Shown in B and C is the average time to first SLE (B) in minutes and average SLE durations (C) in seconds. Shown in D is the carbonyl concentration as a measure of protein peroxidation in brain slices. Shown in E is cell count of PV^+^ interneurons measured through immunostaining. DN – decapitated with nACSF, DA – decapitated with nACSF supplemented with antioxidants, * p<0.05, ** p<0.01, *** p<0.001, **** p<0.0001, n for DN – 7, DA – 6.

### Oxidative damage during brain slice preparation is mediated by hypoxic stress response

Antioxidant supplementation in the cutting solution partially mitigated the loss of inhibitory interneurons and the hyperexcitable network observed with the DN slice preparation method. This observation suggests that the loss of inhibitory interneurons can largely be attributed to oxidative damage during brain slice preparation method. Given that oxidative stress is tightly linked with hypoxia (Merelli et al., 2021), we hypothesize that the hypoxic event during the decapitation procedure leads to this oxidative damage. To examine this hypothesis, we wanted to inhibit the hypoxic response pathway that regulates hypoxic stress response upstream of oxidative stress (Majmundar et al., 2010). This pathway is largely mediated by the hypoxia-inducible factor protein (HIF-1) with the HIF-1α being responsible for acute hypoxia response (Ziello et al., 2007). To modulate this pathway, we supplemented the cutting solution with chrysin, a known inhibitor of HIF-1α (Fu et al., 2007).

Shown in D is the carbonyl concentration as a measure of protein peroxidation in brain slices. Shown in E is cell count of PV^+^ interneurons measured through immunostaining. DN – decapitated with nACSF, DC – decapitated with normal aCSF supplemented with chrysin, * p<0.05, ** p<0.01, *** p<0.001, n for DN – 7, DC – 6.

Addition of chrysin to the cutting solution (see Figure 6) significantly affected the proportion of slices displaying SLEs, *x*^2^ (1, N= 40)=31.3; p<0.0001. Specifically, the proportion of slices displaying SLEs in decapitated mice prepared in nACSF cutting solution containing chrysin (15%) were significantly lower than mice prepared in normal aCSF cutting solution (58.6%). Interestingly, unlike the antioxidant cutting solution, we did not observe any significant difference in the dynamics of the SLE; neither the time to first SLE nor the average SLE durations. Carbonyl concentration in brain slices prepared with cutting solution containing chrysin (DC; 10.4 ± 2.5 nmol/mg protein) was significantly lower than in slices prepared with nACSF cutting solution (DN; 43.0 ± 8.8 nmol/mg protein). Finally, the density of PV^+^ interneurons was similarly significantly higher in slices prepared in cutting solution containing chrysin as compared against slices prepared in nACSF cutting solution (DC, 5.9 ± 1.2 cells/mm^2^; DN, 1.0 ± 0.2 cells/mm^2^)

**Figure 6:**
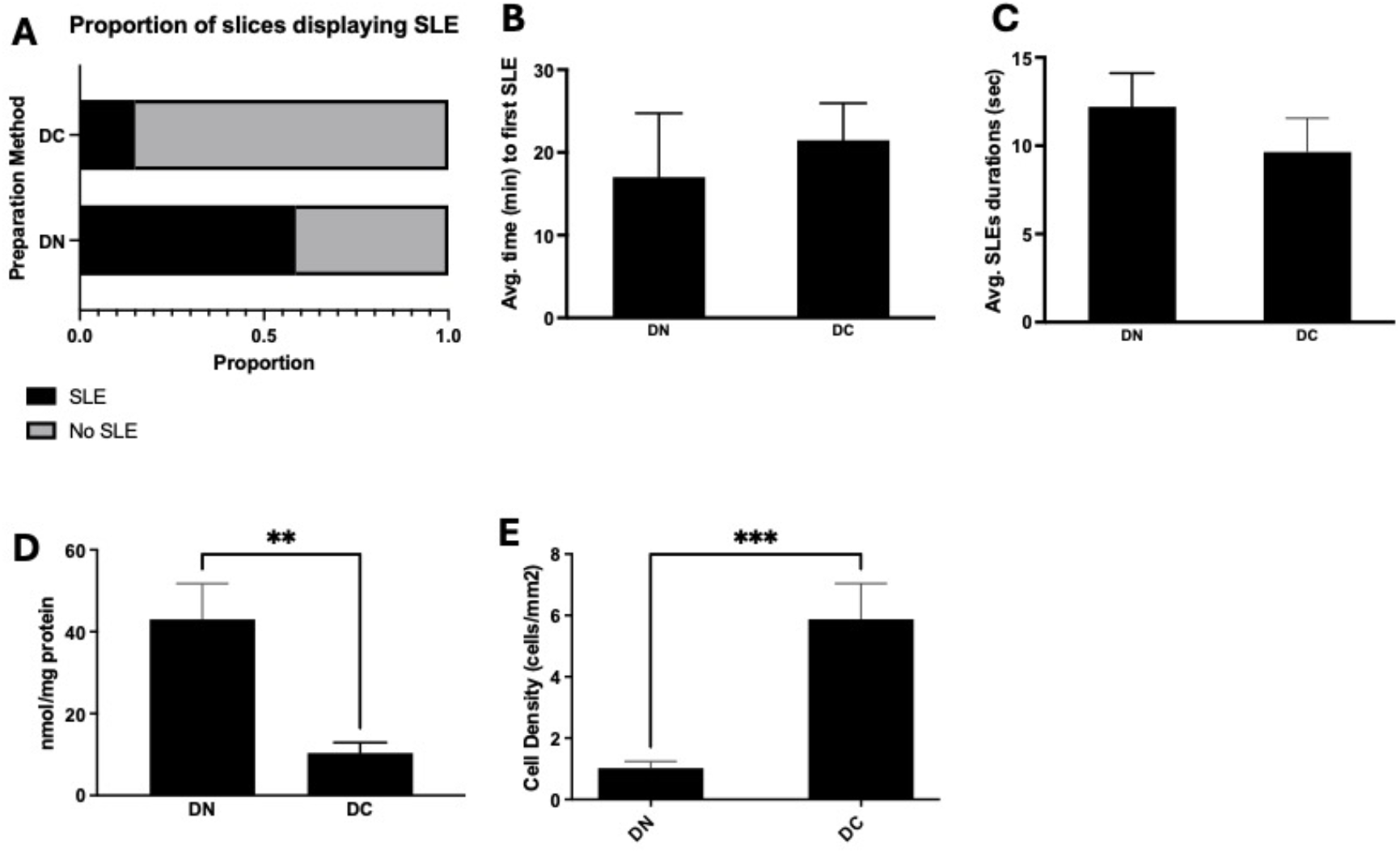
Seizure-like event (SLE) induction is rescued by addition of chrysin, an inhibitor of HIF-1α in the cutting solution. In panel A is shown the proportion of slices displaying SLE in mice prepared with decapitation in nACSF cutting solution supplemented with chrysin (DC) compared against nACSF only cutting solution. Shown in B and C is the average time to first SLE (B) in minutes and average SLE durations (C) in seconds. Shown in D is the carbonyl concentration as a measure of protein peroxidation in brain slices. Shown in E is cell count of PV^+^ interneurons measured through immunostaining. DN – decapitated with nACSF, DC – decapitated with normal aCSF supplemented with chrysin, * p<0.05, ** p<0.01, *** p<0.001, n for DN – 7, DC – 6.

## Discussion

*Ex vivo* acute brain slice electrophysiology has been a mainstay in neuroscience research for the past fifty years and more (Li & McIlwain, 1957) – yet, surprisingly, very few studies have comprehensively characterized the degree of neuronal cell preservation following acute brain slice preparation. We found that brain slice preparation method significantly affected the survival and preservation of neuronal population in the brain slice. In particular, we found that the combination of transcardial perfusion and the use of sACSF preserves the most viable neuronal population. Our current findings are consistent with previous reports showing that sACSF preserves neuronal populations more effectively than nACSF) when used as a cutting solution (Aghajanian & Rasmussen, 1989; Richerson & Messer, 1995). However, interestingly, when brain slices were prepared using cervical dislocation followed by decapitation rather than transcardial perfusion, this difference becomes less apparent. This suggests that the method of using cervical dislocation followed by decapitation (a Schedule 1 method (UK Animals (Scientific Procedures) Act 1986)) induces a baseline level of damage to brain slices, which could not be mitigated using sACSF. In the framework of the 3Rs (Lee et al., 2020), transcardial perfusion with sACSF can be considered a refinement, as it serves as a protective technique that enhances the preservation of brain circuitry, particularly GABAergic interneurons. This experimental refinement is important for critical *in vitro* brain slice physiological studies where neuronal network functions (Kuenzi et al., 2000) are to be examined.

The density of inhibitory and excitatory neuronal populations was diminished during brain slice preparation; however, the use of transcardial perfusion in combination with sACSF significantly mitigated this neuronal loss. We predicted that inhibitory interneurons, in particular PV+ interneurons, could be rapidly lost to oxidative damage during brain slice preparation from its inherent high metabolic nature (Wang & Michaelis, 2010); but we had not anticipated that excitatory neurons would be similarly lost during brain slice preparation. However, cervical dislocation causes rapid respiratory arrest (Carbone et al., 2012) and the subsequent hypoxia-ischemia is known to bring about damage and cell death in cortical pyramidal neurons (Akulinin et al., 1997; Barakat et al., 2024; Sadowski et al., 1999). Markedly increased extracellular glutamate levels during hypoxia (López-Pérez et al., 2012), combined with impaired glutamate reuptake (Vicente et al., 2009), are also likely to contribute to excitotoxic conditions that underlie the significant neurodegeneration of glutamatergic neurons observed. The degeneration of layer III pyramidal neurons in the entorhinal cortex was especially pronounced. This finding may be due to the high levels of recurrent excitatory synaptic connections between these cells and a relatively high degree of electrical coupling (Dhillon & Jones, 2000). Early loss of inhibitory interneuron in neurodegeneration has also been shown to lead to a latter loss of excitatory neurons in the brain (Maestú et al., 2021; Montañana-Rosell et al., 2024; Tweedy et al., 2021), suggesting a bilateral relationship between inhibitory loss and excitatory neurodegeneration. It is interesting to note that in both human and pre-clinical animal models of temporal lobe epilepsy, neurodegeneration in layer III of the mEC is a key feature (Du et al., 1995; Du & Schwarcz, 1992; Du et al., 1993; Kim et al., 1990).

Immunocytochemical analysis of specific inhibitory interneuron subtypes revealed that different brain slice preparation methods preferentially preserved distinct interneuron populations. Using the most effective neuronal preservation method - transcardial perfusion with sACSF - PV^+^ and CB^+^ interneurons showed the highest levels of preservation. This finding was unsurprising given these cell types high metabolic activity (Gulyás et al., 2006) and the vulnerability of PV^+^ interneurons to elevated glutamatergic tone and associated oxidative stress (Behrens et al., 2007; Cantu et al., 2015). NMDAR expression on PV^+^ interneurons also contributes to the selective vulnerability of this cell type in this context. PV^+^ interneurons in the superficial layers of the mEC have a significant synaptic NMDA component (Jones & Bühl, 1993) in comparison to other PV^+^ interneurons in other cortical regions. Excessive glutamate is known to activate NMDARs on PV^+^ interneurons and cause an abnormal influx of calcium ions. This high intracellular calcium burden will in turn lead to neuronal damage/death via a combination of oxidative stress, energy failure and activation of proteolytic enzymes. CB^+^ interneurons have more heterogeneous expression of mitochondrial respiratory enzymes. However, in temporal lobe regions such as the mEC, their mitochondrial expression is as high as parvalbumin-expressing interneurons (Gulyás et al., 2006). Interestingly, decapitation with sACSF specifically preserved VIP^+^ and CCK^+^ interneurons. It is unclear why these specific subclass of interneurons are preserved by decapitation with sACSF; but CCK^+^ and VIP^+^ interneurons are known to have their own circuitry (Nguyen et al., 2020).

Given the significant differences in neuronal populations across various brain slice preparation methods, we next examined the neurophysiological dynamics in *ex vivo* brain slice electrophysiological recordings. Consistently, brain slices prepared with nACSF, particularly when combined with cervical dislocation and decapitation, exhibited characteristics of hyperexcitability. The probability of successfully inducing seizure-like events (SLEs) using an acute epilepsy model was significantly higher in brain slices prepared via cervical dislocation and decapitation. In the case of the 0 Mg^2+^ model a progressive breakdown of GABAergic inhibition plays a crucial role in the evolution of induced epileptic activity (Whittington et al., 1995). This previous finding suggests that for this particular acute seizure model to work a reduction in levels of GABAergic inhibition is required. In the present study, this alteration in inhibition is reflected by the decreased proportion of PS slices generating SLEs and the significant time taken for SLEs to occur in contrast to DN slices. Both these parameters are shown to be linked to the degree of inhibition in the brain as feedforward inhibition has been shown consistently to modulate seizure dynamics (Liou et al., 2018; Trevelyan & Schevon, 2013). Thus, the preservation of inhibitory interneurons in the ‘more viable’ brain slice preparation method has not only affected the success of SLE induction in *in vitro* acute epilepsy model; but also inherently changed the SLEs induced in these brain slices. Thus, yet again, another philosophical question arises whether SLEs induced in conditions of ‘high inhibition’ is similar in nature to SLEs induced in conditions of ‘low inhibition’ in the brain slice.

To identify the mechanism of neuronal loss during brain slice preparation, we devised several rescue experiments. Given the high metabolic rate of parvalbumin-expressing interneurons (Gulyás et al., 2006) and their susceptibility to oxidative damage (Wang & Michaelis, 2010), we hypothesized that the loss of neurons during brain slice preparation is primarily due to oxidative stress. Indeed, application of a cocktail of antioxidants in the nACSF cutting solution significantly attenuated the loss of parvalbumin-expressing interneurons, which subsequently corrected the hyperexcitability traits of these brain slices. Importantly, the SLE dynamics is also more akin to the profile of an sACSF-prepared slices; indicating a potent preservation of inhibition by antioxidant addition to nACSF. Given the period of anoxia experienced by the brain in the decapitation procedure, we suspected that hypoxia response pathway may be upstream of the oxidative stress experienced by the brain slices (Majmundar et al., 2010). To test this, we applied HIF-1α inhibitors in the nACSF cutting solution and while this also rescued parvalbumin-expressing interneurons and the hyperexcitability in the slice, it does not significantly change the profile of the SLEs induced. Collectively, this indicates that oxidative stress is central to the loss of neurons incurred during brain slice preparation and this may be partially regulated by HIF-1α-dependent mechanisms. Likely, there are other factors at-play that collectively contribute to the damage incurred during brain slice preparation such as HIF-2-dependent mechanism (Ratcliffe, 2007), excitotoxicity (Schurr et al., 1995), and mitochondrial dysfunction (Fried et al., 2014).

Regardless, our study has raised important question with wide-ranging implications for neuroscience research. The method by which brain slice preparation is conducted significantly affects the neuronal population preserved in the brain slice and ultimately, the network physiology and dynamics. Thus far, many brain slice studies conducted had not considered the impact of their brain slice preparation method on the interpretations of their results. This is especially important when considering the popular use of *ex vivo* brain slice preparation in acute epilepsy model (Morris et al., 2023), pharmacological studies (Burman et al., 2019), and metabolic studies (Qi et al., 2021). Given our results that indicate a higher likelihood of generating epileptiform activity with decapitation model, it is unsurprising that many studies, even to this day, still utilize brain slices prepared with decapitation with nACSF in acute *ex vivo* epilepsy studies (Cerovic et al., 2023; Dong et al., 2020; Gonzalez-Sulser et al., 2011). While the use of this brain slice preparation method offers a reliable success in induction of SLEs, our study raised an important question – whether these SLEs are ‘artificially’ induced by a particularly damaging brain slice preparation method that does not preserve inhibition. This is especially interesting as some of these brain slice models are lauded as *in vitro* drug-resistant epilepsy model (Burman et al., 2019; Cerovic et al., 2023) but given the lack of preservation of physiological inhibition in these brain slices, the lack of pharmacological response of some of these anticonvulsant drugs would need to be re-evaluated with this consideration in mind.

While this study comprehensively evaluated the metabolic changes in brain slice preparation, there are factors that we had not examined in the current study. A recent study found that microglia and inflammation is significantly affected by different brain slice preparation method (Berki et al., 2024). Given the significant metabolic change with different brain slice preparation method, we suspected that glial cells would be significantly affected as well in our models, and this could be an important future direction for our study. Another crucial factor in brain slice preparation is the temperature of the cutting solution. In our current study, we have maintained this variable at ice-cold temperature, a popular choice in brain slice preparation; but some studies have shown that using cutting solutions at physiological temperature also significantly changed the neuronal dynamics in the brain slice (Eguchi et al., 2020; Huang & Uusisaari, 2013); arguably closer to a physiological state. Temperature significantly changes an organ metabolic state and thus, this is again another variable that would be interesting to explore in future studies.

## Conclusion

Our study is the first to comprehensively characterize the impact of different brain slice preparation method on the preservation of neuronal population and their network dynamics in the resultant brain slices. We found that the combination of transcardial perfusion of rodents and the use of sucrose-based artificial cerebrospinal fluid as cutting solution is the best brain slice preparation method to preserve inhibition in the brain slice. Arguably, this preservation of inhibition is a crucial goal for brain slice preparation to model physiological brain state. The loss of inhibitory interneurons incurred in brain slice preparation that leads to this loss of inhibition is due to oxidative stress experienced during brain slice preparation. Limiting this oxidative stress during brain slice preparation can preserve inhibition in the brain and should be considered as neuroprotective strategy during *ex vivo* brain slice preparation. This has a broad impact for the use of *ex vivo* brain slice preparation for epilepsy studies and future studies using this popular model should consider the impact that the brain slice preparation has on the physiological and metabolic state of the tissue.

## List of Abbreviations

4-AP: 4-aminopyridine
ACSF: artificial cerebrospinal fluid
DA: decapitation with normal artificial cerebrospinal fluid with antioxidants
DC: decapitation with normal artificial cerebrospinal fluid with chrysin
DN: decapitation with normal artificial cerebrospinal fluid
DNPH: 2-dinitrophenylhydrazine
DS: decapitation with sucrose artificial cerebrospinal fluid
GABA: gamma-aminobutyric acid
HIF: hypoxia-inducible factor
LRD: late-recurrent discharge
LTP: long term potentiation
nACSF: normal artificial cerebrospinal fluid
NMDA-R: N-methyl-D-aspartate receptor
PN: perfused with normal artificial cerebrospinal fluid
PS: perfused with sucrose artificial cerebrospinal fluid
sACSF: sucrose artificial cerebrospinal fluid
SLE: seizure-like event

## Data Availability Statement –

All datasets generated and analyzed during this study are available from the corresponding principal investigators upon reasonable request.

## References

Aghajanian, G. K., & Rasmussen, K. (1989). Intracellular studies in the facial nucleus illustrating a simple new method for obtaining viable motoneurons in adult rat brain slices. Synapse, 3(4), 331–338. 10.1002/syn.890030406

Akulinin, V. A., Stepanov, S. S., Semchenko, V. V., & Belichenko, P. V. (1997). Dendritic changes of the pyramidal neurons in layer V of sensory-motor cortex of the rat brain during the postresuscitation period. Resuscitation, 35(2), 157–164. 10.1016/S0300-9572(97)00048-8

Anderson, W. W., Anderson, W. W., Lewis, D. V., Scott Swartzwelder, H., & Wilson, W. A. (1986). Magnesium-free medium activates seizure-like events in the rat hippocampal slice. Brain Research, 398(1), 215–219. 10.1016/0006-8993(86)91274-6

Avegno, E. M., Middleton, J. W., & Gilpin, N. W. (2019). Synaptic GABAergic transmission in the central amygdala (CeA) of rats depends on slice preparation and recording conditions. Physiological Reports, 7(19), e14245. 10.14814/phy2.14245

Barakat, R. M., Turcani, M., Al-Khaledi, G., Kilarkaje, N., Al-Sarraf, H., Sayed, Z., & Redzic, Z. (2024). Low oxygen in inspired air causes severe cerebrocortical hypoxia and cell death in the cerebral cortex of awake rats. Neuroscience Letters, 818, 137515. 10.1016/j.neulet.2023.137515

Barbarosie, M., & Avoli, M. (1997). CA3-driven hippocampal-entorhinal loop controls rather than sustains in vitro limbic seizures. J Neurosci, 17(23), 9308–9314. 10.1523/jneurosci.17-23-09308.1997

Barbarosie, M., Louvel, J., D’Antuono, M., Kurcewicz, I., & Avoli, M. (2002). Masking synchronous GABA-mediated potentials controls limbic seizures. Epilepsia, 43(12), 1469–1479. 10.1046/j.1528-1157.2002.17402.x

Behrens, M. M., Ali, S. S., Dao, D. N., Lucero, J., Shekhtman, G., Quick, K. L., & Dugan, L. L. (2007). Ketamine-induced loss of phenotype of fast-spiking interneurons is mediated by NADPH-oxidase. Science, 318(5856), 1645–1647. 10.1126/science.1148045

Berki, P., Cserép, C., Környei, Z., Pósfai, B., Szabadits, E., Domonkos, A., Kellermayer, A., Nyerges, M., Wei, X., Mody, I., Kunihiko, A., Beck, H., Kaikai, H., Ya, W., Lénárt, N., Wu, Z., Jing, M., Li, Y., Gulyás, A. I., & Dénes, Á. (2024). Microglia contribute to neuronal synchrony despite endogenous ATP-related phenotypic transformation in acute mouse brain slices. Nature Communications, 15(1), 5402. 10.1038/s41467-024-49773-1

Brückner, C., & Heinemann, U. (2000). Effects of standard anticonvulsant drugs on different patterns of epileptiform discharges induced by 4-aminopyridine in combined entorhinal cortex–hippocampal slices. Brain Research, 859(1), 15–20. 10.1016/S0006-8993(99)02348-3

Burman, R. J., Selfe, J. S., Lee, J. H., van den Berg, M., Calin, A., Codadu, N. K., Wright, R., Newey, S. E., Parrish, R. R., Katz, A. A., Wilmshurst, J. M., Akerman, C. J., Trevelyan, A. J., & Raimondo, J. V. (2019). Excitatory GABAergic signalling is associated with benzodiazepine resistance in status epilepticus. Brain, 142(11), 3482–3501. 10.1093/brain/awz283

Cantu, D., Walker, K., Andresen, L., Taylor-Weiner, A., Hampton, D., Tesco, G., & Dulla, C. G. (2015). Traumatic Brain Injury Increases Cortical Glutamate Network Activity by Compromising GABAergic Control. Cereb Cortex, 25(8), 2306–2320. 10.1093/cercor/bhu041

Carbone, L., Carbone, E. T., Yi, E. M., Bauer, D. B., Lindstrom, K. A., Parker, J. M., Austin, J. A., Seo, Y., Gandhi, A. D., & Wilkerson, J. D. (2012). Assessing cervical dislocation as a humane euthanasia method in mice. J Am Assoc Lab Anim Sci, 51(3), 352–356.

Cerovic, M., Di Nunzio, M., Craparotta, I., & Vezzani, A. (2023). An in vitro model of drug-resistant seizures for selecting clinically effective antiseizure medications in Febrile Infection-Related Epilepsy Syndrome [Original Research]. Frontiers in Neurology, Volume 14 - 2023. 10.3389/fneur.2023.1129138

Chan, F., Lax, N. Z., Voss, C. M., Aldana, B. I., Whyte, S., Jenkins, A., Nicholson, C., Nichols, S., Tilley, E., Powell, Z., Waagepetersen, H. S., Davies, C. H., Turnbull, D. M., & Cunningham, M. O. (2019). The role of astrocytes in seizure generation: insights from a novel in vitro seizure model based on mitochondrial dysfunction. Brain, 142(2), 391–411. 10.1093/brain/awy320

Dhillon, A., & Jones, R. S. G. (2000). Laminar differences in recurrent excitatory transmission in the rat entorhinal cortex in vitro. Neuroscience, 99(3), 413–422. 10.1016/S0306-4522(00)00225-6

Dong, L., Li, G., Gao, Y., Lin, L., Zhang, K.-h., Tian, C.-x., Cao, X.-b., & Zheng, Y. (2020). Effect of priming low-frequency magnetic fields on zero-Mg2+-induced epileptiform discharges in rat hippocampal slices. Epilepsy Research, 167, 106464. 10.1016/j.eplepsyres.2020.106464

Du, F., Eid, T., Lothman, E. W., Köhler, C., & Schwarcz, R. (1995). Preferential neuronal loss in layer III of the medial entorhinal cortex in rat models of temporal lobe epilepsy. J Neurosci, 15(10), 6301–6313. 10.1523/jneurosci.15-10-06301.1995

Du, F., & Schwarcz, R. (1992). Aminooxyacetic acid causes selective neuronal loss in layer III of the rat medial entorhinal cortex. Neurosci Lett, 147(2), 185–188. 10.1016/0304-3940(92)90591-t

Du, F., Whetsell, W. O., Jr., Abou-Khalil, B., Blumenkopf, B., Lothman, E. W., & Schwarcz, R. (1993). Preferential neuronal loss in layer III of the entorhinal cortex in patients with temporal lobe epilepsy. Epilepsy Res, 16(3), 223–233. 10.1016/0920-1211(93)90083-j

Eguchi, K., Velicky, P., Hollergschwandtner, E., Itakura, M., Fukazawa, Y., Danzl, J. G., & Shigemoto, R. (2020). Advantages of Acute Brain Slices Prepared at Physiological Temperature in the Characterization of Synaptic Functions [Methods]. Frontiers in Cellular Neuroscience, Volume 14 - 2020. 10.3389/fncel.2020.00063

Ezeriņa, D., Takano, Y., Hanaoka, K., Urano, Y., & Dick, T. P. (2018). N-Acetyl Cysteine Functions as a Fast-Acting Antioxidant by Triggering Intracellular H(2)S and Sulfane Sulfur Production. Cell Chem Biol, 25(4), 447–459.e444. 10.1016/j.chembiol.2018.01.011

Fekete, A., & Wang, L.-Y. (2025). Spotting multifaced actions of magnesium on NMDA receptors. Neuron, 113(7), 963–965. 10.1016/j.neuron.2025.03.011

Fried, N. T., Moffat, C., Seifert, E. L., & Oshinsky, M. L. (2014). Functional mitochondrial analysis in acute brain sections from adult rats reveals mitochondrial dysfunction in a rat model of migraine. American Journal of Physiology-Cell Physiology, 307(11), C1017–C1030. 10.1152/ajpcell.00332.2013

Fu, B., Xue, J., Li, Z., Shi, X., Jiang, B. H., & Fang, J. (2007). Chrysin inhibits expression of hypoxia-inducible factor-1alpha through reducing hypoxia-inducible factor-1alpha stability and inhibiting its protein synthesis. Mol Cancer Ther, 6(1), 220–226. 10.1158/1535-7163.Mct-06-0526

Gęgotek, A., & Skrzydlewska, E. (2022). Antioxidative and Anti-Inflammatory Activity of Ascorbic Acid. Antioxidants (Basel), 11(10). 10.3390/antiox11101993

Gonzalez-Sulser, A., Wang, J., Motamedi, G. K., Avoli, M., Vicini, S., & Dzakpasu, R. (2011). The 4-aminopyridine in vitro epilepsy model analyzed with a perforated multi-electrode array. Neuropharmacology, 60(7), 1142–1153. 10.1016/j.neuropharm.2010.10.007

Gulyás, A. I., Buzsáki, G., Freund, T. F., & Hirase, H. (2006). Populations of hippocampal inhibitory neurons express different levels of cytochrome c. European Journal of Neuroscience, 23(10), 2581–2594. 10.1111/j.1460-9568.2006.04814.x

Hashimoto, A., Sawada, T., & Natsume, K. (2017). The change of picrotoxin-induced epileptiform discharges to the beta oscillation by carbachol in rat hippocampal slices. Biophys Physicobiol, 14, 137–146. 10.2142/biophysico.14.0_137

Heuzeroth, H., Wawra, M., Fidzinski, P., Dag, R., & Holtkamp, M. (2019). The 4-Aminopyridine Model of Acute Seizures in vitro Elucidates Efficacy of New Antiepileptic Drugs. Front Neurosci, 13, 677. 10.3389/fnins.2019.00677

Huang, S., & Uusisaari, M. Y. (2013). Physiological temperature during brain slicing enhances the quality of acute slice preparations [Methods]. Frontiers in Cellular Neuroscience, Volume 7 - 2013. 10.3389/fncel.2013.00048

Jones, R. S. (1989). Ictal epileptiform events induced by removal of extracellular magnesium in slices of entorhinal cortex are blocked by baclofen. Exp Neurol, 104(2), 155–161. 10.1016/s0014-4886(89)80009-3

Jones, R. S., da Silva, A. B., Whittaker, R. G., Woodhall, G. L., & Cunningham, M. O. (2016). Human brain slices for epilepsy research: Pitfalls, solutions and future challenges. J Neurosci Methods, 260, 221–232. 10.1016/j.jneumeth.2015.09.021

Jones, R. S., Heinemann, U. F., & Lambert, J. D. (1992). The entorhinal cortex and generation of seizure activity: studies of normal synaptic transmission and epileptogenesis in vitro. Epilepsy Res Suppl, 8, 173–180. 10.1016/b978-0-444-89710-7.50027-6

Jones, R. S. G., & Bühl, E. H. (1993). Basket-like interneurones in layer II of the entorhinal cortex exhibit a powerful NMDA-mediated synaptic excitation. Neuroscience Letters, 149(1), 35–39. 10.1016/0304-3940(93)90341-H

Kageyama, G. H., & Wong-Riley, M. T. T. (1982). Histochemical localization of cytochrome oxidase in the hippocampus: Correlation with specific neuronal types and afferent pathways. Neuroscience, 7(10), 2337–2361. 10.1016/0306-4522(82)90199-3

Kim, J. H., Guimaraes, P. O., Shen, M. Y., Masukawa, L. M., & Spencer, D. D. (1990). Hippocampal neuronal density in temporal lobe epilepsy with and without gliomas. Acta Neuropathol, 80(1), 41–45. 10.1007/bf00294220

Kuenzi, F. M., Fitzjohn, S. M., Morton, R. A., Collingridge, G. L., & Seabrook, G. R. (2000). Reduced long-term potentiation in hippocampal slices prepared using sucrose-based artificial cerebrospinal fluid. Journal of Neuroscience Methods, 100(1), 117–122. 10.1016/S0165-0270(00)00239-9

Lee, K. H., Lee, D. W., & Kang, B. C. (2020). The ‘R’ principles in laboratory animal experiments. Laboratory Animal Research, 36(1), 45. 10.1186/s42826-020-00078-6

Levine, R. L., Garland, D., Oliver, C. N., Amici, A., Climent, I., Lenz, A.-G., Ahn, B.-W., Shaltiel, S., & Stadtman, E. R. (1990). [49] Determination of carbonyl content in oxidatively modified proteins. In Methods in Enzymology (Vol. 186, pp. 464–478). Academic Press. 10.1016/0076-6879(90)86141-H

Li, C.-L., & McIlwain, H. (1957). Maintenance of resting membrane potentials in slices of mammalian cerebral cortex and other tissues in vitro. The Journal of Physiology, 139(2), 178–190. 10.1113/jphysiol1957.sp005885

Li Zhang, C., Dreier, J. P., & Heinemann, U. (1995). Paroxysmal epileptiform discharges in temporal lobe slices after prolonged exposure to low magnesium are resistant to clinically used anticonvulsants. Epilepsy Research, 20(2), 105–111. 10.1016/0920-1211(94)00067-7

Lim, L., Mi, D., Llorca, A., & MarÍn, O. (2018). Development and Functional Diversification of Cortical Interneurons. Neuron, 100(2), 294–313. 10.1016/j.neuron.2018.10.009

Liou, J. Y., Ma, H., Wenzel, M., Zhao, M., Baird-Daniel, E., Smith, E. H., Daniel, A., Emerson, R., Yuste, R., Schwartz, T. H., & Schevon, C. A. (2018). Role of inhibitory control in modulating focal seizure spread. Brain, 141(7), 2083–2097. 10.1093/brain/awy116

López-Pérez, S. J., Morales-Villagrán, A., Ventura-Valenzuela, J., & Medina-Ceja, L. (2012). Short- and long-term changes in extracellular glutamate and acetylcholine concentrations in the rat hippocampus following hypoxia. Neurochemistry International, 61(2), 258–265. 10.1016/j.neuint.2012.03.009

Maestú, F., de Haan, W., Busche, M. A., & DeFelipe, J. (2021). Neuronal excitation/inhibition imbalance: core element of a translational perspective on Alzheimer pathophysiology. Ageing Research Reviews, 69, 101372. 10.1016/j.arr.2021.101372

Majmundar, A. J., Wong, W. J., & Simon, M. C. (2010). Hypoxia-inducible factors and the response to hypoxic stress. Mol Cell, 40(2), 294–309. 10.1016/j.molcel.2010.09.022

Merelli, A., Repetto, M., Lazarowski, A., & Auzmendi, J. (2021). Hypoxia, Oxidative Stress, and Inflammation: Three Faces of Neurodegenerative Diseases. J Alzheimers Dis, 82(s1), S109–s126. 10.3233/jad-201074

Mody, I., Lambert, J. D., & Heinemann, U. (1987). Low extracellular magnesium induces epileptiform activity and spreading depression in rat hippocampal slices. Journal of Neurophysiology, 57(3), 869–888. 10.1152/jn.1987.57.3.869

Montañana-Rosell, R., Selvan, R., Hernández-Varas, P., Kaminski, J. M., Sidhu, S. K., Ahlmark, D. B., Kiehn, O., & Allodi, I. (2024). Spinal inhibitory neurons degenerate before motor neurons and excitatory neurons in a mouse model of ALS. Science Advances, 10(22), eadk3229. 10.1126/sciadv.adk3229

Morris, G., Avoli, M., Bernard, C., Connor, K., de Curtis, M., Dulla, C. G., Jefferys, J. G. R., Psarropoulou, C., Staley, K. J., & Cunningham, M. O. (2023). Can in vitro studies aid in the development and use of antiseizure therapies? A report of the ILAE/AES Joint Translational Task Force. Epilepsia, 64(10), 2571–2585. 10.1111/epi.17744

Muldoon, S. F., Villette, V., Tressard, T., Malvache, A., Reichinnek, S., Bartolomei, F., & Cossart, R. (2015). GABAergic inhibition shapes interictal dynamics in awake epileptic mice. Brain, 138(10), 2875–2890. 10.1093/brain/awv227

Müller, S., Guli, X., Hey, J., Einsle, A., Pfanz, D., Sudmann, V., Kirschstein, T., & Köhling, R. (2018). Acute epileptiform activity induced by gabazine involves proteasomal rather than lysosomal degradation of KCa2.2 channels. Neurobiology of Disease, 112, 79–84. 10.1016/j.nbd.2018.01.005

Nguyen, R., Venkatesan, S., Binko, M., Bang, J. Y., Cajanding, J. D., Briggs, C., Sargin, D., Imayoshi, I., Lambe, E. K., & Kim, J. C. (2020). Cholecystokinin-Expressing Interneurons of the Medial Prefrontal Cortex Mediate Working Memory Retrieval. J Neurosci, 40(11), 2314–2331. 10.1523/jneurosci.1919-19.2020

Papouin, T., & Haydon, P. G. (2018). Obtaining Acute Brain Slices. Bio Protoc, 8(2). 10.21769/BioProtoc.2699

Qi, G., Mi, Y., & Yin, F. (2021). Characterizing brain metabolic function ex vivo with acute mouse slice punches. STAR Protoc, 2(2), 100559. 10.1016/j.xpro.2021.100559

Raimondo, J. V., Heinemann, U., de Curtis, M., Goodkin, H. P., Dulla, C. G., Janigro, D., Ikeda, A., Lin, C.-C. K., Jiruska, P., Galanopoulou, A. S., & Bernard, C. (2017). Methodological standards for in vitro models of epilepsy and epileptic seizures. A TASK1-WG4 report of the AES/ILAE Translational Task Force of the ILAE. Epilepsia, 58(S4), 40–52. 10.1111/epi.13901

Ratcliffe, P. J. (2007). HIF-1 and HIF-2: working alone or together in hypoxia? J Clin Invest, 117(4), 862–865. 10.1172/jci31750

Richerson, G. B., & Messer, C. (1995). Effect of composition of experimental solutions on neuronal survival during rat brain slicing. Exp Neurol, 131(1), 133–143. 10.1016/0014-4886(95)90015-2

Robeson, C. D., & Baxter, J. G. (1943). α-Tocopherol, a Natural Antioxidant in a Fish Liver Oil. Journal of the American Chemical Society, 65(5), 940–943. 10.1021/ja01245a047

Sadowski, M., Wisniewski, H. M., Jakubowska-Sadowska, K., Tarnawski, M., Lazarewicz, J. W., & Mossakowski, M. J. (1999). Pattern of neuronal loss in the rat hippocampus following experimental cardiac arrest-induced ischemia. Journal of the Neurological Sciences, 168(1), 13–20. 10.1016/S0022-510X(99)00159-8

Samoilova, M., Li, J., Pelletier, M. R., Wentlandt, K., Adamchik, Y., Naus, C. C., & Carlen, P. L. (2003). Epileptiform activity in hippocampal slice cultures exposed chronically to bicuculline: increased gap junctional function and expression. Journal of Neurochemistry, 86(3), 687–699. 10.1046/j.1471-4159.2003.01893.x

Schurr, A., Payne, R. S., Heine, M. F., & Rigor, B. M. (1995). Hypoxia, excitotoxicity, and neuroprotection in the hippocampal slice preparation. J Neurosci Methods, 59(1), 129–138. 10.1016/0165-0270(94)00203-s

Ting, J. T., Kalmbach, B., Chong, P., de Frates, R., Keene, C. D., Gwinn, R. P., Cobbs, C., Ko, A. L., Ojemann, J. G., Ellenbogen, R. G., Koch, C., & Lein, E. (2018). A robust ex vivo experimental platform for molecular-genetic dissection of adult human neocortical cell types and circuits. Sci Rep, 8(1), 8407. 10.1038/s41598-018-26803-9

Trevelyan, A. J., & Schevon, C. A. (2013). How inhibition influences seizure propagation. Neuropharmacology, 69, 45–54. 10.1016/j.neuropharm.2012.06.015

Tweedy, C., Kindred, N., Curry, J., Williams, C., Taylor, J. P., Atkinson, P., Randall, F., Erskine, D., Morris, C. M., Reeve, A. K., Clowry, G. J., & LeBeau, F. E. N. (2021). Hippocampal network hyperexcitability in young transgenic mice expressing human mutant alpha-synuclein. Neurobiol Dis, 149, 105226. 10.1016/j.nbd.2020.105226

Vicente, É., Degerone, D., Bohn, L., Scornavaca, F., Pimentel, A., Leite, M. C., Swarowsky, A., Rodrigues, L., Nardin, P., Vieira de Almeida, L. M., Gottfried, C., Souza, D. O., Netto, C. A., & Gonçalves, C. A. (2009). Astroglial and cognitive effects of chronic cerebral hypoperfusion in the rat. Brain Research, 1251, 204–212. 10.1016/j.brainres.2008.11.032

Wang, X., & Michaelis, E. (2010). Selective neuronal vulnerability to oxidative stress in the brain [Review]. Frontiers in Aging Neuroscience, 2. 10.3389/fnagi.2010.00012

Whittington, M. A., Traub, R. D., & Jefferys, J. G. (1995). Erosion of inhibition contributes to the progression of low magnesium bursts in rat hippocampal slices. J Physiol, 486 (Pt 3)(Pt 3), 723–734. 10.1113/jphysiol1995.sp020848

Wu, J., Cai, Y., Wu, X., Ying, Y., Tai, Y., & He, M. (2021). Transcardiac Perfusion of the Mouse for Brain Tissue Dissection and Fixation. Bio Protoc, 11(5), e3988. 10.21769/BioProtoc.3988

Ziello, J. E., Jovin, I. S., & Huang, Y. (2007). Hypoxia-Inducible Factor (HIF)-1 regulatory pathway and its potential for therapeutic intervention in malignancy and ischemia. Yale J Biol Med, 80(2), 51–60.

